# Structured neural fluctuations can generate noise invariance and inter-areal gating at distinct timescales

**DOI:** 10.1101/2024.07.05.602210

**Authors:** Jorrit S. Montijn, J. Alexander Heimel

## Abstract

The brain processes, computes, and categorizes sensory input. But even in sensory brain areas, the relationship between input signals and neuronal spiking activity is complex and non-linear. Fast subsecond fluctuations in neuronal population responses dominate the temporal dynamics of neural circuits. Traditional approaches have treated this activity as “noise” that can be averaged away by taking the mean spiking rate over wide time bins or over multiple trial repetitions, but this ignores much of the temporal dynamics that naturally occur in neural systems. We find that subsecond flares of increased population activity are layer– and cell-type specific, and large-scale computational modelling suggests they may serve as an inter-areal gating mechanism. Moreover, we find that most of the neural variability is restricted to a population-gain axis. This observation explains why neural systems can function in the presence of excessive variability: population-level spiking dynamics generate invariance to the majority of neural noise.

## Introduction

Sensory stimuli evoke reproducible patterns of neuronal responses in the brain^1^. The reliability of these patterns, or neural codes, influences how fast and accurate animals can respond to sensory stimuli^2–7^. Paradoxically, however, neural codes show a large degree of variability^8–15^. This has led many to ask: how does noise impact neural codes? Studies in this highly active topic of research have revealed that not all types of noise are equally detrimental^16–19^. In recent years, experimental and theoretical work has shown that the interaction between noise and neuronal population codes can be best understood using a geometric framework of neural manifolds^20–23^. In many instances, this theory provides a more parsimonious explanation of how neural activity affects behavioural output than could be formulated using a single-neuron perspective^24–26^. This framework has contributed to many theoretical advances, but makes the simplifying assumption that neural codes are stationary. Recent work has challenged this assumption by showing that neural codes drift over the time course of days^27–34^. Whether neural codes are stationary at short timescales is not yet known, but investigating this is of critical importance if we wish to build an accurate model of how neural circuits compute.

Animals can make quick decisions within a hundred milliseconds^35–38^, so neural computations must therefore occur within a fraction of these response latencies. And yet, most experimental studies on neural codes and computations investigate timescales one or two orders of magnitude longer, while many theoretical studies ignore temporal dynamics altogether^17,19^. Work on the membrane voltage dynamics of single cells has shown that neurons go through up-states (and down-states), where they are more (or less) likely to generate action potentials for a given stimulus^39–43^. Whether these single-cell dynamics translate into population-level phenomena is currently unknown. If populations of neurons show synchronized dynamics, this would suggest that over the timecourse of a single trial, population codes may not be stable either. It is therefore of critical importance to understand how subsecond dynamics of neuronal population spiking activity interact with neural coding, and link these findings to the more thoroughly studied field of neural coding at the timescale of trial averages.

In this study, we investigated the temporal dynamics of neuronal population activity using Neuropixels recordings in the primary visual cortex of mice, in combination with large-scale computational modelling of the early visual system. We found that at a subsecond timescale, neuronal population activity was excessively variable – it was characterized by intermittent flares of activity on a quiescent background. These flares were cortical-layer specific and cell-type specific. Computational modelling showed they can appear without the need for feedback connections and can generate feedforward subcircuit-switching or “gating”. Secondly, we found that the temporal dynamics of neuronal population activity were highly structured from a geometric perspective. This prompted us to develop a new model for neural noise where the variability is predominantly aligned to a population “gain axis”. Compared to the predictive power of a multivariate log-normal distribution (R^2^ = 0.517), our gain model performed exceptionally well (R^2^ = 0.994), despite using only half the number of parameters. Based on these results, we propose an updated formulation for neuronal population codes that incorporates this gain-axis as a scaling variable to which the feature encoding is invariant. In our updated, more parsimonious model for neural codes, manifolds therefore become gain-scalable entities, where the bulk of neural noise is restricted to a population gain axis that does not reduce neural information. We therefore conclude that the majority of the seemingly excessive variability of neuronal population activity is not noise. It is highly structured to avoid interfering with neural codes, and likely reflects ongoing neural computations, such as divisive normalization and the initiation of subcircuit switching of downstream areas.

## Results

### Flares in spiking activity dominate neural dynamics at a subsecond timescale

We recorded neuronal spiking activity in the primary visual cortex (V1) of awake mice using Neuropixels, while presenting drifting gratings (Fig. 1a,b). A classical approach in neuroscience is to average neural activity over trial repetitions (Fig. 1c,d). This gives an impression of a rather stable response during a trial in terms of spiking rate and stimulus encoding reliability, perhaps with the exception of the initial onset response peak (Fig. 1e,f). This stability, however, is to a large degree an artefact of averaging. When zooming in on the population response during a single trial, it appears highly variable, marked by occasional flares of strong activity flanked by deep troughs of quiescent periods (Fig. 1g). These short-term fluctuations are often assumed to be random noise, and may be a necessary property of having binary signals like spiking events, which by definition cannot produce a continuous signal.

**Figure 1.**
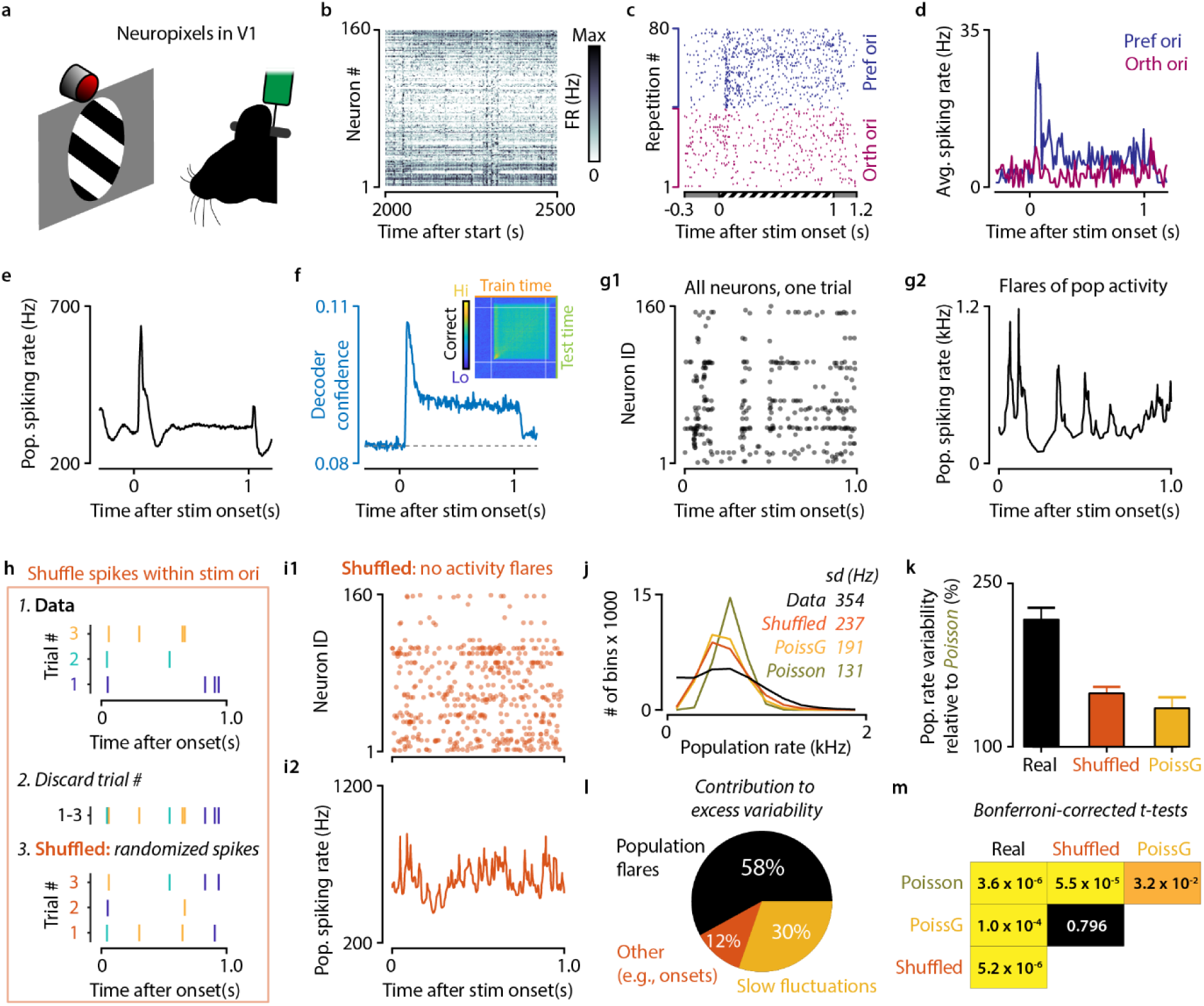
Short timescale dynamics of neuronal populations are characterized by flares of neural activity. a) Schematic of experimental design, showing recordings with Neuropixels in mouse V1 during drifting grating presentation. b) Long-timescale example of population activity, showing mostly stable patterns. c,d) Spike raster plot and peri-stimulus time histogram of example neuron in response to its preferred and orthogonal stimulus. e,f) Averaging over trials and neurons shows a stable population response (e) and stimulus information (f). g) Spike-raster plot (left) and instantaneous population spiking rate (right) during a single trial, showing flares of spiking activity on a relatively quiescent background. h) A spike-shuffling procedure designed to remove population flares, but keep all other properties of spiking dynamics intact. For each neuron and stimulus class, spikes are reassigned to random trials, preserving the number of spikes per trial and latency per spike. At a trial-averaged level, shuffled data is therefore indistinguishable from unshuffled data. i) As g, but for shuffled data, showing the absence of population flares. j) Distribution of population spiking rates in 33 ms bins for an example recording. Real data are more variable than shuffled data (orange), gain-modulated Poisson process cells (yellow), and standard spiking-rate matched Poisson process cells (olive green). Note that real data show an overabundance of bins with low firing rates (quiescent periods) as well as high firing rates (flares). k) Variability normalized per recording to rate-matched Poisson process cells (mean +/− SEM of sd over bins for n = 11 recordings). l) A large majority of excess variability in population spiking can be explained by the presence of flares (58%), a third by slow 1-s fluctuations (30%), and the remainder by other processes, such as response peaks after stimulus onset (12%). m) Real data was significantly more variable than all Shuffled (p=5.2 x 10^−6^), Poisson x Gain (1.0 x 10^−4^), and Poisson data (3.6 x 10^−6^). Shuffled and Poisson x Gain data were more variable than Poisson data (p= 5.5 x 10^−5^ and p=3.2 x 10^−2^ respectively), but did not differ significantly (p=0.796).

The first question we wished to answer was whether these flares of activity could be a simple stochastic side-effect of the intrinsic variability of single neurons. We therefore devised a shuffling procedure that we applied to the data, where we destroyed the temporal coordination between neurons, but kept all other properties intact, including average single neuron firing rates, trial-level spike counts and the temporal structure of the trial, such as onset peaks (Fig. 1h). Destroying the temporal coordination between cells indeed removed much of the short timescale fluctuations, and population activity flares were greatly reduced (Fig. 1i). We also compared the variability of the real and shuffled data to a null-hypothesis that all neurons were spiking as independent Poisson-process cells. We computed the variability of population spiking as the standard deviation (sd) over 33 ms bins (Fig. 1j). Indeed, the real data were more variable than shuffled data, which in turn were more variable than Poisson data (sd over bins, real data = 354 Hz, Shuffled = 237 Hz, Poisson = 131 Hz).

To what extent do longer-timescale fluctuations explain this excess variability? Poisson processes are stationary, but it is known that changes in brain state, such as arousal, can strongly modulate firing rates in V1. These brain states vary at the timescale of seconds, so we generated another data set to investigate the role of these slow fluctuations. Like before, we generated spikes using rate-matched Poisson cells, but modulated each neuron’s firing rate over time according to the population rate in the real data averaged over one second bins. This “Poisson x Gain” data showed an intermediate population variability (sd = 191 Hz), well below that of the real data. We repeated these analyses on all data sets (n = 11 recordings) and normalized the spiking-rate standard deviations to that of the rate-matched Poisson process data for each recording. This insured variability is comparable across recordings, as differences in intrinsic spiking rates and number of recorded neurons are corrected for.

We found that, relative to Poisson process neurons, real data was 116% more variable, shuffled data was 48% more variable, and gain-modulated Poisson-process data was 34% more variable (fig. 1k-m; p = 3.6 x 10^−6^, p = 5.5 x 10^−5^ and p = 3.2 x 10^−2^ respectively). This means that a large majority (1 – 48/116 = 58%) of all excess variability is due to short-term spiking synchronization between neurons (fig. 1k,l). Of the remaining 42% of excess variability, 30% can be explained by longer-timescale fluctuations present in both shuffled and gain-modulated data. Finally, 12% is caused by other processes, such as an onset response, which is present in shuffled data, but not in gain-modulated Poisson data. Therefore, we can conclude that single-trial neural dynamics are dominated by the intermittent appearance of population flares on a quiescent background, and that the influence of these flares strongly exceeds that of slower second-scale fluctuations in population spiking rates.

### Flares are supragranular, densely populated and show constant neural coding efficiency

Our previous analysis showed that real neural activity is much more variable than would be expected by chance, but it also showed that flares of activity are not discrete, stereotyped events. Rather, high-rate and low-rate epochs exist on a continuum, and flares seem to be the result of the higher-end rate epochs. This fluctuating and heterogeneous nature of population flares makes them difficult to study directly, but we still wished to investigate the possible mechanisms and functions of this exceptionally wide range of population activity. We therefore designed an analysis based on splitting the population data into blocks of an equal number of spikes (fig. 2a,b). One can compare it to a peri-stimulus histogram, where a variable number of spikes is put into a bin with a fixed duration (e.g., 33 ms). Instead in our spike-block analysis, we fixed the number of spikes, while making the bin duration variable. This allows us to compare high-rate regimes (flares) with low-rate regimes (quiescent epochs) while having an internal control for many biases as each block contains the exact same number of spikes.

**Figure 2.**
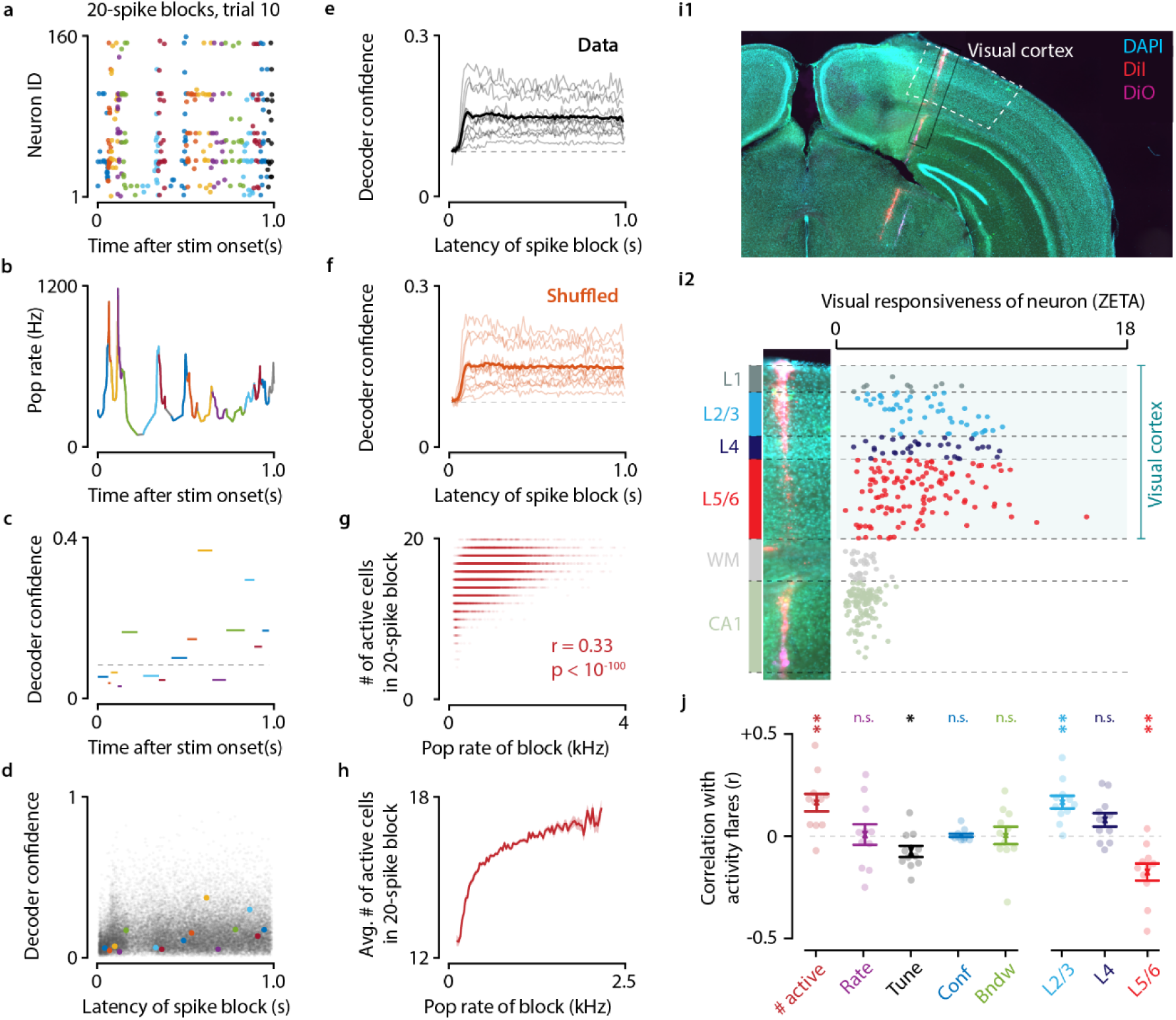
Flares show decreased population sparseness and originate primarily from supragranular cells. a,b) Spike-raster plot (a) and instantaneous population firing rate (b) of the same trial as depicted in Figure 1i1. Each colour denotes a block of 20 consecutive spikes. c) Each spike block can be used as a “trial” to train a decoder, yielding an estimate of the neural information (decoder confidence) for each spike block. d) As c, showing the spike blocks of a-c as coloured dots on a background of all spike blocks of one example recording. e,f) Using spike block decoding, the onset peak in stimulus information disappears (compare with Figure 1f), which shows that the amount of information per spike is roughly constant over the entire trial duration. g) For each spike block, we counted the number of participating cells to investigate if population sparseness differs between flares and quiescent periods. This scatter plot shows a strong, positive correlation with the average population rate of the block (Pearson r = 0.30, p<10^−100^). h) Same as g, but data are now averaged over bins with 25 Hz steps. i) Histological reconstruction of the Neuropixels probe trajectories in combination with electrophysiological markers allowed us to very accurately determine the resident cortical layer for each recorded unit. j) A spike-block based analysis of the correlation between a block’s average population firing rate and various properties showed that population flares are associated with an increase in number of participating cells (red, p = 8.0 x 10^−3^), a small reduction in average orientation tuning (black, p = 0.026), an overrepresentation of supragranular cells (light blue, p = 2.1 x 10^−3^), and an underrepresentation of infragranular cells (light red, p = 4.9 x 10^−3^).

Using this analysis, we decoded the stimulus orientation using a fixed number of 20 spikes and found a strikingly constant level of information over the course of a trial (fig. 2c-f). Peaks in information during onsets (e.g., fig. 1f) therefore seem to be a side-effect from simply having more spikes in one time bin rather than from an increase in coding efficiency. Now that we designed a way of interrogating how various aspects of neural spiking dynamics differ between low-rate and high-rate regimes, we next investigated the population sparseness. For each block, we counted the number of distinct neurons contributing at least one spike, and plotted this as a function of the population rate during the block (fig. 2g,h). We found a strong positive correlation (Pearson r = 0.33, p < 10^−100^, n = 26793 spike blocks). This means that during flares of neural activity, the population response is less sparse: more different neurons are active, while during troughs of activity, the baseline rate is carried by smaller sets of cells.

Cells in supragranular layers (L2/3) tend to show sparser responses than in infragranular layers (L5/6)^44–46^, so we wondered if the difference in population sparseness could be explained by a layer-specific role in generating population flares. We therefore split the data into supragranular, granular, and infragranular cells (fig. 2i), and repeated the above analysis, but now computed the fraction of (supra/infra-)granular cells during each spike block. Indeed, as we suspected, we found a strong layer-specific effect: L2/3 cells were overrepresented during flares (p = 2.3 x 10^−3^), while L5/6 cells were underrepresented (p = 4.9 x 10^−3^), and L4 cells showed no effect (p = 0.060) (fig. 2j).

We also investigated several properties related to the neural coding of orientation. This included the average firing rate of cells contributing at least 1 spike to the block, the orientation tuning strength of these cells, and their bandwidth. Interestingly, of these three properties only the average orientation tuning strength seemed somewhat lower during flares than quiescent periods (p = 0.026), whereas the average firing rate and bandwidth were uncorrelated (p > 0.98). This general lack of correlation with coding-related properties was reflected in the decoder’s confidence: this was equally high during flares and quiescent periods (p = 0.99). This suggests that the neural code is equally efficient in terms of information per spike during peaks of activity as during troughs. These results suggest that the function of flares – if any – is not related to the fidelity of the neural code in V1.

### Large-scale simulations suggest an inter-areal gating function for V1 flares

The previous results suggest that flares originate primarily from supragranular layers, which are known to project to feedforward cortical areas. We therefore hypothesized that flares might function as feedforward gating events that increase the spiking probability of specific postsynaptic subcircuits. Since this hypothesis is difficult to test *in vivo*, we instead used a large-scale model of the early visual system we developed previously^47^, where we could more easily interrogate inter-areal interactions (fig. 3a). This model simulates an LGN stage where each neuron spikes stochastically with a probability proportional to the stimulation of either an on-center or off-center filter applied to an input image. These spikes are then sent to a V1 layer, where all cells are simulated as leaky integrate-and-fire neurons. V1 is sparsely recurrently connected, and excitatory cells in V1 project to V2 cells, which are themselves also recurrently connected. For the following analyses we created 11 pseudo populations of model neurons, one for each Neuropixels session, and matched the number of model neurons to the number of neurons in the real data.

**Figure 3.**
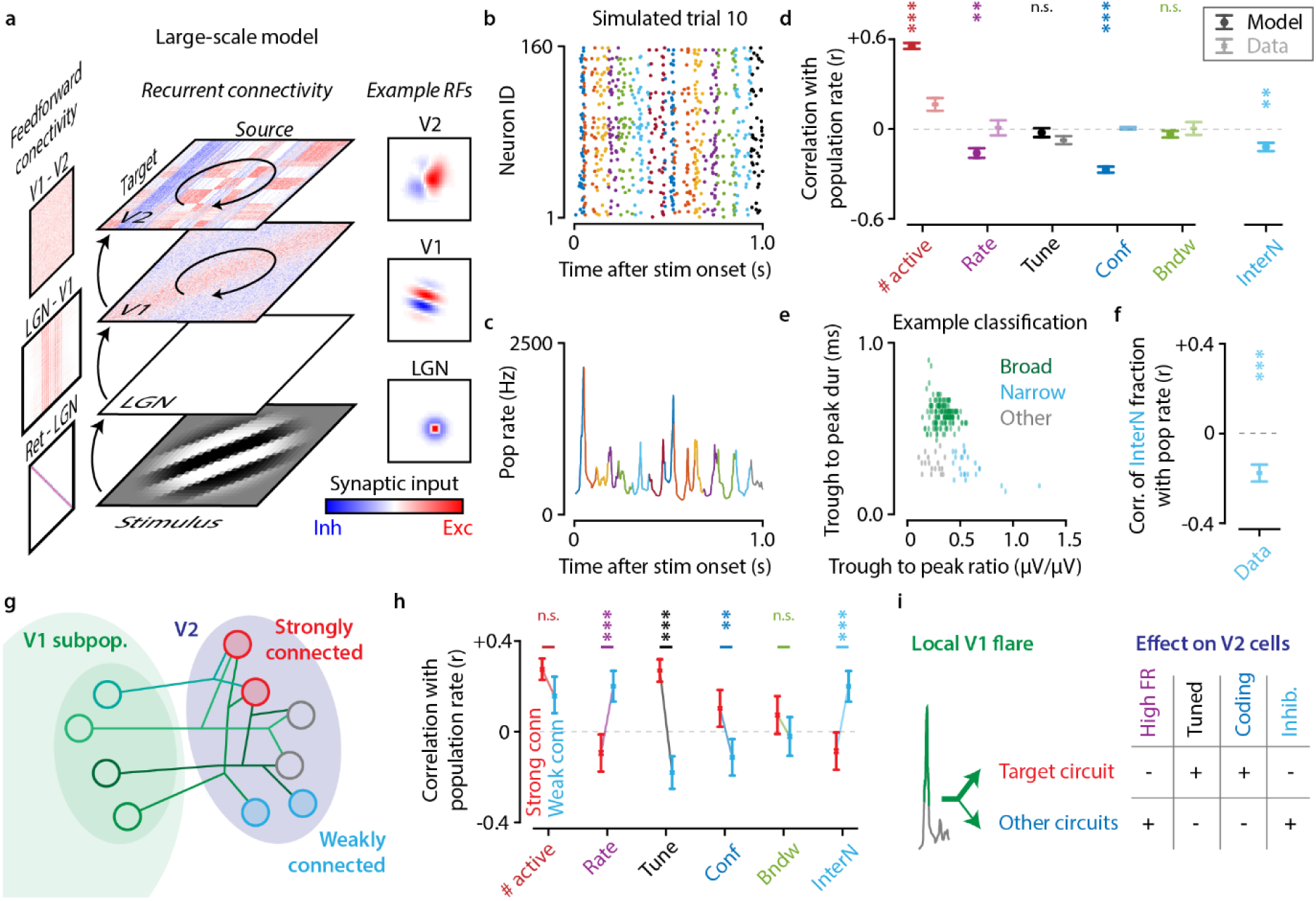
Large-scale simulations suggest a gating function for population flares. a) Schematic of our feedforward computational model of the early visual system with recurrent intra-areal connectivity. b,c) Spike raster plot and instantaneous population rate of an example trial, showing the occurrence of population flares in simulated data. d) Similar to flares in real data, simulated flares are associated with an increase in number of participating cells (red, p = 4.1 x 10^−10^). The model also shows a reduction in average cell firing rate (purple, p = 1.1 x 10^−3^) and a surprisingly strong reduction in decoder confidence (blue, p = 2.4 x 10^−7^). In the model the fraction of interneurons in active cells is also reduced during flares (light blue, p = 2.4 x 10^−3^), suggesting a shift in excitatory/inhibitory balance may play role in the formation of flares. e) Single units were classified by waveform properties into broad spiking (green), narrow spiking (blue), and other (grey) cells. f) We used this classification to investigate the involvement of interneurons (narrow waveform units) in population flares. Similar to simulated data, the excitation/inhibition ratio was strongly correlated with population activity rate (p = 1.6 x 10^−4^). g) We hypothesized that V1 flares might act as an inter-areal gating mechanism, and affect V2 spiking responses differently between strongly and weakly connected cells in the target area. We therefore split the simulated V2 cells into strongly (red), weakly (blue), and intermediately connected cells. h) We found that V1 flares have a differential effect on V2 subcircuits: within the strongly connected V2 circuit, cells with low firing rates (purple) and strong stimulus tuning (black) became more active, while interneuron activity was reduced (blue), leading to an increase in decoder confidence (blue). In contrast, in weakly connected V2 circuits, the opposite was true: primarily cells with high firing rates (purple) and weak tuning (black) were active during flares, and interneuron activity was enhanced (blue), leading to a decrease in decoder confidence (blue). i) These results suggest that spatiotemporally localized V1 flares may activate specific, strongly connected V2 subcircuits, while suppressing other subcircuits.

We found that this simple feedforward model reproduces the population flares we found in real neural data, although the quiescent periods in the model appeared less prominent (fig. 3b,c). Still, the model shows that flare-like dynamics can easily arise from simple (recurrent) connectivity structures, and that flares can be generated locally without the need for feedback connections or fluctuating modulatory input. We repeated the aforementioned spike-block analysis on these model data, and found largely similar results. Like in the real data, high-activity epochs showed more distributed neuron involvement (fig. 3d, # active, p = 4.1 x 10^−10^), but we did observe a reduction in decoder confidence that was not present in the neural data (fig. 3d, Conf, p = 2.4 x 10^−7^). This difference may in part be explained by the generally higher decoder confidence and accuracy in the model data, which widened the range of confidence values and thereby increases the statistical power to observe changes in the confidence.

We also observed that the balance of active cells during high-rate epochs shifts to be more dominated by excitatory cells (fig. 3d, InterN, p = 2.4 x 10^−3^). This suggests that, in the model, local interneurons may be involved in the generation of population flares. We tested this hypothesis in the real data by classifying all units by their waveform properties into broad-spiking, narrow-spiking and undefined/intermediate cells^48^ (fig. 3e). We repeated the analysis as above, and found that also in the real data, there was a highly significant shift in excitation/inhibition balance between flares and quiescent periods (fig. 3f, p = 1.6 x 10^−4^).

Having confirmed that our model reproduces the most important aspects of the temporal dynamics we observed in real neural circuits, we turned to answering our hypothesis that flares may function as a feedforward gating mechanism. For each pseudo-recording (n=11), we then selected two V2 subpopulations of the same size (on average 5.3% of 1200 V2 cells), one that contained the V2 cells most strongly connected to the V1 subpopulation, and one that contained the V2 cells most weakly connected to the V1 subpopulation (fig. 3g). We defined the connection strength for each V2 cell as the average over synaptic weights between the cell and the V1 subpopulation. We then separately analysed both V2 subpopulations using spike-blocks, where for each spike-block we calculated the population rate of V1 cells. We then computed the correlation of the various spike-block metrics with the V1 rate during that spike-block. This way, we could test how the strength of V1 activity influences V2 activity, and whether this differed between V2 subpopulations that were strongly or weakly connected to the V1 subpopulation.

We reproduced two previous within-V1 observations: during strong V1 activity, the V2 population shows more distributed activity, in both the strongly-connected and weakly-connected subpopulations (weak vs strong, p = 0.059) (fig. 3h). Secondly, there was no effect on the bandwidth of active V2 cells (p = 0.265). All other properties showed strong cross-over effects. During V1 flares, within the strongly-connected V2 subpopulation, the activated V2 cells have a lower firing rate and stronger tuning, the decoder confidence is increased, and the inhibitory/excitatory balance is reduced. In contrast, within the weakly-connected population, we found the opposite: activated V2 cells have a higher firing rate and weaker tuning, the decoder confidence is decreased, and the inhibitory/excitatory balance is increased. These differences in correlation with V1 activity between weakly– and strongly-connected V2 subpopulations were highly significant (Rate, p = 9.1 x 10^−5^; Tune, p = 2.8 x 10^−13^; Conf, p = 7.5 x 10^−3^; InterN, p = 1.6 x 10^−4^). The strongly differentiated response within V2 subcircuits in the model, in combination with the layer-specific V1 involvement we found in real V1 data, confirms that one function of V1 flares may indeed be to differentially activate and suppress specific subcircuits in postsynaptic populations (fig. 3i). That said, this remains a theoretically plausible hypothesis for now, until future studies can perform a more direct experiment to confirm – or falsify – these results *in vivo*.

### Different timescales show different dominant neural dynamics

So far we have focused our attention on short timescale fluctuations. We have seen that neural dynamics at the subsecond scale are dominated by intermittent flares and quiescent periods, but how does this relate to the more thoroughly studied dynamics of single trials that occur on the order of seconds? Longer timescales are less variable, as variability in population rates averages out over longer binning windows (fig. 4a). This explains why flares have not been reported previously in studies on inter-trial variability, but the question remains at what timescale the effects of flares start to average out. To link the short-timescale variability of flares to the wider literature on neural variability, we therefore investigated how neural variability changes as a function of timescale, and which properties in the neural dynamics are responsible.

**Figure 4.**
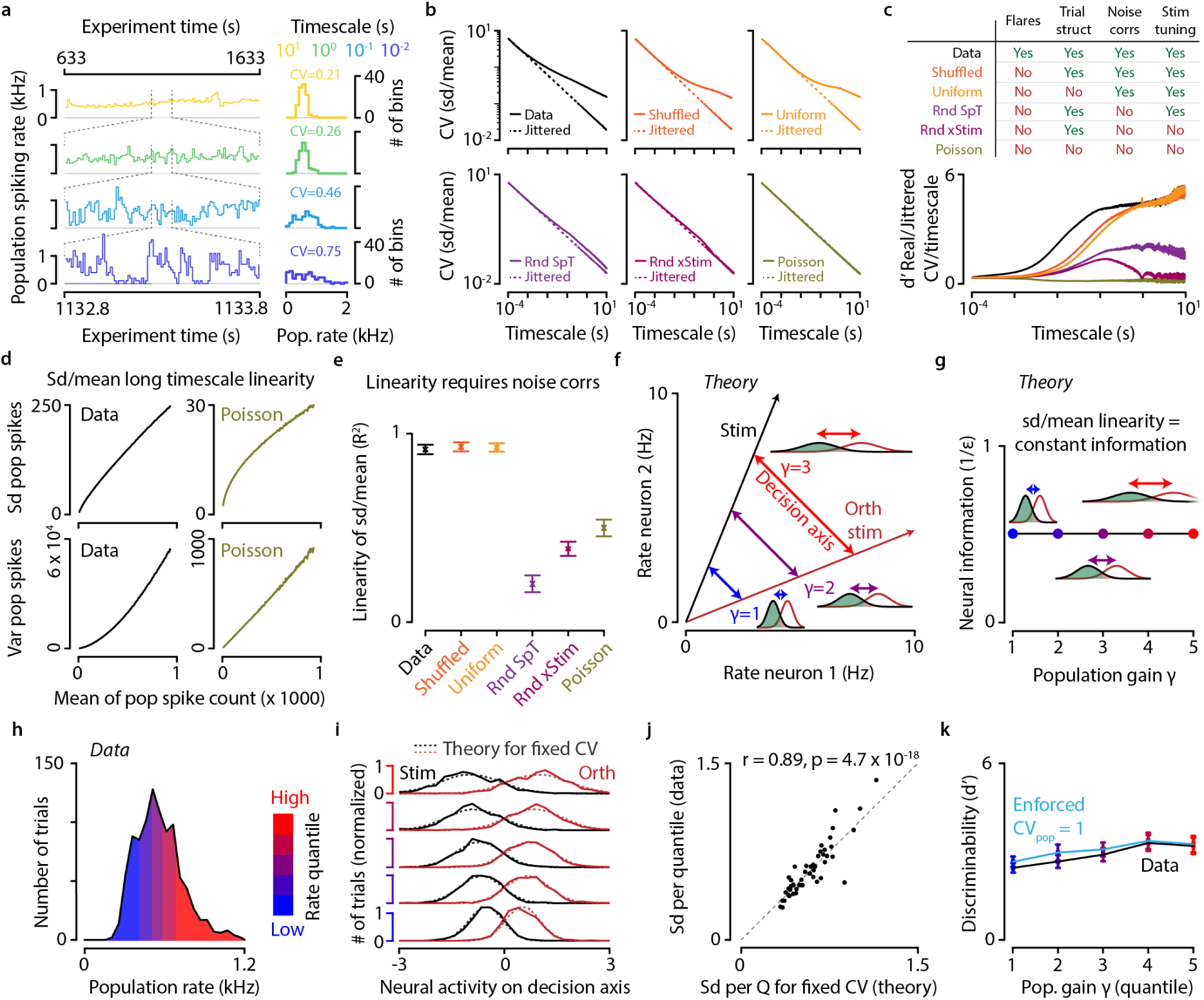
Flares dominate neural dynamics at short timescales, whereas a linear neural scaling law emerges at longer timescales. a) Example of neural variability at timescales varying from 10 ms to 10 s. Short timescales are intrinsically more variable. b,c) Neural data can be randomized in different ways to disentangle the effects of flares, stimulus onset, trial-level noise correlations, and neuronal tuning to different stimuli. b) Coefficient of variation (CV) as a function of timescale for real data (Data, black); shuffled data (Shuffled, orange); uniformized within-trial spike latency data (Uniform, yellow); randomized within-stimulus class spike time data (Rnd SpT, purple); randomized across-stimulus class spike time data (Rng xStim, red); and random Poisson process data (Poisson, olive green). c) Flares cause an excess in spiking variability across timescales of 1 – 100 ms, but at 100 – 1000 ms timescales there is no difference between data with flares (black) and without flares (Shuffled, Uniform). In contrast, at timescales of 1 – 10 s, trial-level noise correlations (present in Data, Shuffled, and Uniform) explain over half of super-Poissonian variability. The remaining 40% of excess variability at long timescales is caused by neuronal responses differing between stimulus classes (Rnd SpT vs. Rnd xStim and Poisson). d) At longer timescales, real data is approximately linear in standard deviation (sd) over the mean population spike count per bin (top left). In contrast, Poisson process neurons cause population spike counts to be linear in variance (sd^2^) over mean instead. Data show an example recording. e) The sd/mean near-linearity present at longer timescales is dependent on trial-level noise correlations (Data, Shuffled, Uniform), as removal of these noise correlations leads to considerably lower estimates of sd/mean linearity (Rnd SpT, Rnd xStim, Poisson). Mean ± SEM over recordings (n=11). f) Example theoretical schematic showing sd/linearity over the stimulus class decision axis for different levels of population gain γ (blue, purple, red). g) If the sd/mean ratio is constant over a range of population gains, then neural information is also constant. h) Distribution of population firing rate per trial for an example recording split into five quantiles of 192 trials each. i) Projecting the population activity for each activity quantile onto the decision axis reveals that real neural data shows a near-linear relationship between mean and standard deviation. Solid lines: data. Dotted lines: theory for an exactly linear relationship. j) Quantification of (i), showing the standard deviation (across trials) in neural activity projected onto the decision axis for an exactly constant sd/mean (x-axis) and real data (y-axis). Each dot is one quantile of one recording. k) Mean +/− SEM over recordings, showing the discriminability of orthogonal stimulus orientations for all trials split by population activity quantile. Real data (black) matches well with surrogate data where sd/mean linearity is mathematically enforced (blue).

This question can be answered using a fairly straightforward approach. For each recording, we binned the spikes of all neurons over the entire recording using various binning widths, ranging from 0.1 ms up to 10 s. At each timescale, we compute the coefficient of variation (CV); the standard deviation of the number of spikes per bin divided by the mean, and compare this CV to what we obtain if we jitter all spikes uniformly across the entire recording (fig. 4b). This way, we can correct for the intrinsic change in variability as a function of timescale, and use the difference between the real and jittered curve as a metric for the amount of supra-Poisson variability at any timescale. Performing this analysis on real data, we find that neural dynamics are supra-Poisson at timescales longer than 1-10 ms. We repeated this analysis for different randomizations of the data, which we designed to remove specific processes to interrogate their effect on generating neural variability (fig. 4c). First, we used Shuffled data, as described above, where only flares are removed, but the data still contains the trial structure, inter-trial noise correlations, and stimulus tuning. We found that, as expected, long-timescale variability is not affected, but Shuffled data shows a marked decrease in variability (compared to real data) between roughly 1 – 100 ms. This means that the flares/quiescent epochs we observed previously dominate neural dynamics specifically at this range around 10 ms. At timescales longer than 100 ms, flares average out, and other processes take over.

We then repeated the above analysis on data where we jittered spikes within single trials (“Uniform”), destroying both flares and trial structure, to investigate whether the variability we observed around 100 ms timescales was dependent on a stereotyped stimulus-onset response. Surprisingly, we found barely any difference with Shuffled data (fig. 4c), indicating that the variability we found at longer timescales can be almost entirely explained by trial-level variability (noise correlations). To test this further, we removed trial-level noise correlations by shuffling spike times across stimulus repetitions (“Rnd SpT”), and indeed found a 60% decrease in variability at longer timescales (fig. 4c). Variability was decreased, but not to the extent that it was indistinguishable from completely jittered data. We hypothesized that the remaining 40% of supra-Poisson variability may be caused by different stimulus orientations evoking a different neural response. We therefore repeated the analysis with a randomization where we also destroyed the stimulus information (“Rnd xStim”) and found that this indeed brought the long timescale variability down to the level of a stationary Poisson process (“Poisson”). We can therefore conclude that flares dominate neural dynamics at timescales between 1 – 100 ms. They average out over longer timescales, where noise correlations and stimulus-specific responsiveness together generate slower fluctuations that can be seen at the level of single trials.

### A linear neural scaling law emerges at longer timescales

We were intrigued that the effect of population flares dissipates at longer timescales, but is replaced by a similarly strong supra-Poisson variability of a different nature. At short timescales, we observed a surprisingly constant level of neural information, despite exceptionally variable spiking rates. We therefore wondered if a similar constancy might be found at longer timescales. We calculated the standard deviation (sd), variance and mean spike count for the data at timescales longer than 100 ms, and found that while the real data is almost linear in sd/mean, a Poisson process is instead linear in variance/mean (fig. 4d). We quantified this linearity by the explained variance (R^2^) of a line fitted through the sd/mean points. We repeated this analysis for all recordings and data randomizations, we found a stark difference in linearity between data that preserved trial-level noise correlations (R^2^ for Data = 0.92, Shuffled = 0.93, Uniform = 0.93) and those where they were destroyed (R^2^ for Rnd SpT = 0.20, Rnd xStim = 0.39, Poisson = 0.50).

This sd/mean linearity hints at an interesting possibility: as we will explain below, if the sd/mean linearity holds across the decision axis of the stimulus tuning manifold, then this implies that the neural information is constant, regardless of the gain of population activity (fig. 4f,g). We also provide a simple supporting proof for this in the methods. We further investigated whether this constancy in information holds in our data. For each recording, we calculated the population rate for each trial and split the trials into five quantiles, from low to high rates (fig. 4h). For each subdivision, we trained logistic decoders to decode all stimulus orientations from their orthogonal orientation. We then projected the population response of all trials onto the decoder’s decision axis. For each activation quantile, we therefore obtain a mean response and standard deviation for both the stimulus and its orthogonal pair (fig. 4i). Now we compare whether the sd of trial activations on the decision axis scales with the distance in means between the two stimulus classes. Indeed, when plotting the empirical sd as a function of the theoretical expectation when the sd/mean relationship were fixed, we found a very strong correlation (fig. 4j; Pearson r = 0.89, p = 4.7 x 10^−18^). As a final test we also calculated the stimulus discriminability (d’) for the data, as a function of population gain quantiles, and compared this with modified data where we enforced a coefficient of variation (CV = sd/mean) of exactly one. The data were again remarkably similar (fig. 4k). These results show that neuronal population activity at a trial-level scales in a very particular way: the variability grows linearly with the population gain, which leads to a roughly constant level of stimulus information.

### Noise in neuronal population responses is predominantly gain-aligned

Research on neural coding, variability and noise correlations (including our own), has commonly used a fairly normative framework: the neural code can be approximated by averaging neuronal responses over trials, and any trial-level deviation from this neural code, or manifold, is noise. Whether this noise impairs neural coding depends on its geometric structure, since only variability along the tangent of a manifold will lead to misclassification^18,27,47^. The standard neural noise model used to study these topics is the multivariate Gaussian, but our previous results suggest that neural noise may contain more structure than is captured by a Gaussian model. Specifically, the observation that neural information is constant, regardless of the gain of population activity (fig. 4f-k), led us to hypothesize that the population gain may define a specific type of noise.

We compared this baseline model of Gaussian noise (fig. 5a) with our alternative hypothesis that neural noise is predominantly aligned along a gain-axis (fig. 5b). In the normal model we estimated *2n* parameters for each recording (*n* being the number of neurons): the mean and standard deviation of each neuron’s spiking rate over trials (1 second duration). For our gain model, we used only *n + 2* parameters. The gain axis requires *n* parameters and is defined by the average rate of each neuron. The other two parameters define the mean and standard deviation of a log-normal distribution over this gain axis. All parameters were computed in closed-form from the neural data and therefore required no fitting. We quantified the performance of each model by sorting the neural responses of all trials and generating the same number of (sorted) samples from the noise model. Plotting the generated values as a function of the real data is therefore a version of a Q-Q (quantile-quantile) plot.

**Figure 5.**
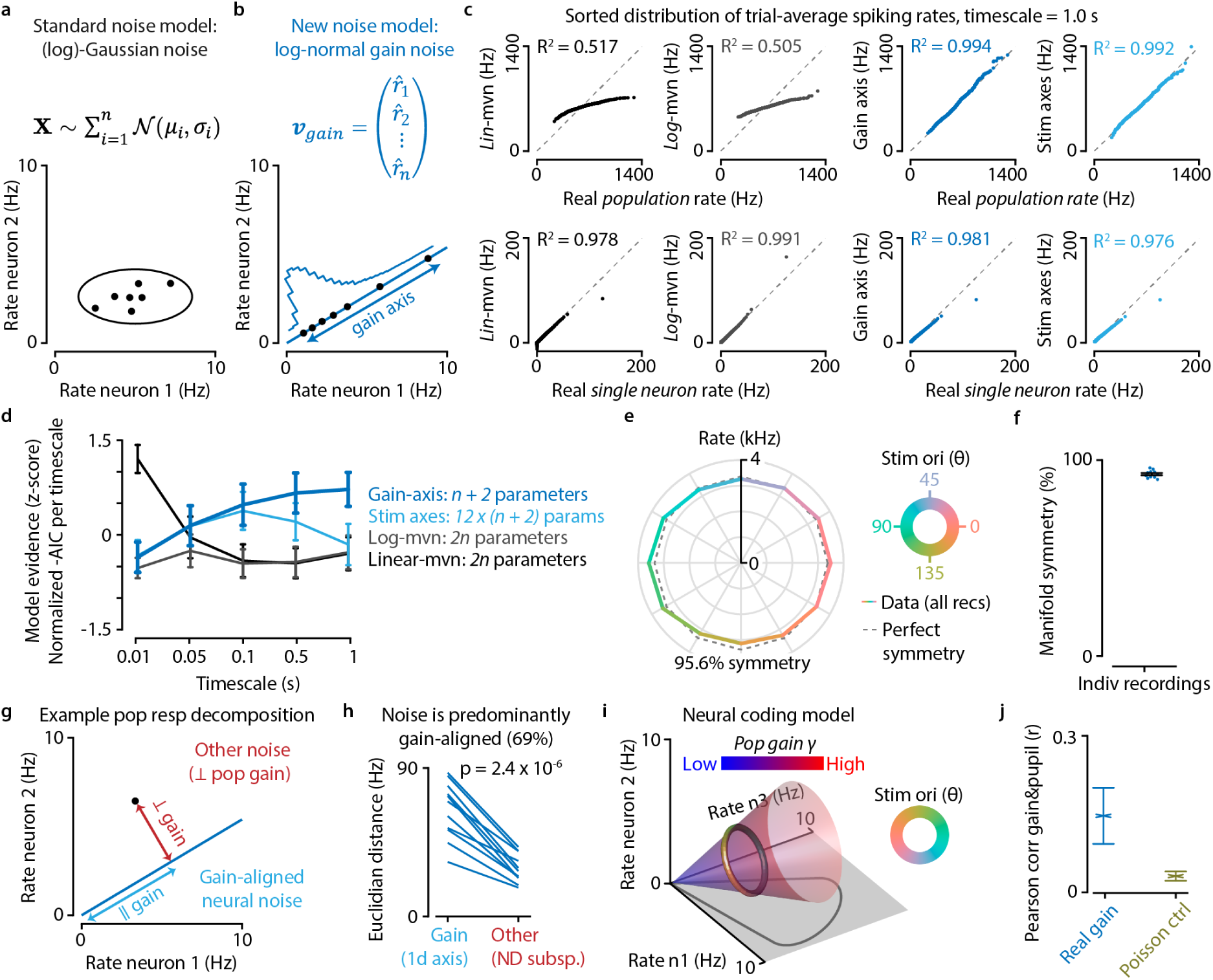
A population-gain based perspective on neural coding offers a more parsimonious model of neural activity. a,b) Alternative theoretical frameworks to describe neural noise. a) The standard model for noise in neuronal population activity is a multivariate (log-)normal distribution, where each dot represents a single trial. b) Our alternative model describes neural noise as a log-normal distribution over the population gain axis. c) Example recording showing how well four models describe the distribution of neuronal spiking rates at a timescale of 1 second (one stimulus presentation). Top row: total population activity is described reasonably well by a linear multivariate normal (Lin-mvn; R^2^ = 0.517) and similarly well by a logarithmic multivariate normal (log-mvn; R^2^ = 0.505), but exceptionally well by our log-normal gain model (Gain axis, R^2^= 0.994; stim axes, R^2^ = 0.992). Bottom row: single-neuron responses are described similarly well by all models (R^2^ = 0.976 – 0.991). d) A model-comparison analysis using the Akaike information criterion (AIC) shows our gain model is a more parsimonious description of neuronal population activity at longer timescales (100 ms and above). e) The neural manifold for orientation-tuning is highly symmetric (constant Euclidian vector norm) at 95.6%. f) Individual recordings show symmetries of 90-96%. g) Schematic showing the decomposition of a population response into a gain-aligned (blue) and orthogonal (red) component for a simplified 2D case. h) Neural noise was predominantly aligned along the 1-dimensional gain-axis (69% of neural variability), despite the other directions spanning a much larger 23 – 159 dimensional subspace. i) We propose a model where the neural code for orientation consists of two subspaces: a linear subspace of population gain and a circular dimension of stimulus orientation orthogonal to the gain subspace. This orthogonality allows neural codes to be unaffected by the dominant gain-aligned mode of neural noise. j) Per-trial population gain shows a positive correlation with pupil size (Pearson r = 0.15, p = 2.9 x 10^−2^).

We found that the normal model did a reasonable job of describing the distribution of population firing rates (fig. 5c, *lin-mvn*, R^2^ = 0.517), but predicted a narrower range of population firing rates than we observed in real data. On the other hand, it did an excellent job of describing the distribution of all single-neuron single-trial spiking rates (R^2^ = 0.978). This suggests that single neuron activity is approximately Gaussian, but the assumption of independence leads to reduced variance when averaging across individually noisy neurons. We also observed that the linear normal model predicted some negative values, and it has been previously shown that neuronal spiking rates are better described by a log-normal distribution. We therefore repeated the above analysis for a log-mvn model, but found a very similar performance (*log-mvn*, population R^2^ = 0.505, single-neuron R^2^ = 0.991).

We repeated the analysis for our gain-noise model, where all activity is by definition restricted to a single gain axis. Despite being a much more restrictive rank-1 model with half the number of parameters, it was able to describe the distribution of population and single-neuron firing rates with stunning accuracy (fig. 5c, *gain axis*, population R^2^ = 0.994; single neuron R^2^ = 0.981). While room for further improvement is limited with such a high R^2^, we also investigated if using a separate gain axis for each stimulus class could further improve the model’s predictions, but these predictions were indistinguishable from the single-axis model (*stim axes*, R^2^ = 0.992 and R^2^ = 0.976). We repeated these analyses for all recordings and computed the normalized model evidence (Akaike information criterion) for these four models at different timescales, from 10 ms up to 1 second (single trial). Matching our previous analyses, we found our gain model showed the highest model evidence for timescales of 100 ms and longer (fig. 5d). These results show that neural activity at longer timescales varies predominantly along a population gain-axis.

### A gain-invariant neural coding model

The sd/mean linearity along feature coding dimensions (see Figure 4) in combination with a much stronger gain-dominant neural noise (fig. 5a-d) warrants an updated perspective on neural coding. Note that the following description is for orientation coding specifically, but the same would apply to any other encoded variable. We hypothesize that the neural code can be formulated to consist of two distinct parts: a linear subspace of population gain, and a second orthogonal subspace that contains the stimulus orientation. This orthogonality means that neural codes are unaffected by the dominant gain-aligned mode of noise: the population gain simply becomes a normalizable constant that multiplicatively scales all neural responses (eq 12-16). Information-limiting noise exists, but scales with the population gain (fig. 4h-k). Under this formulation, neural information is invariant to the population gain, so it may be more parsimonious to describe a neural manifold as a gain-scalable entity rather than as a mean-defined entity with a large manifold-orthogonal noise component in the direction of the population gain axis.

One further requirement for this reformulation is that the feature manifold is symmetric, in the sense that it lays in a subspace that is equidistant to the origin. If this were not the case, then a difference in orientation would be encoded in part as a difference in the population gain, and the feature manifold and gain subspace would not be orthogonal. We therefore computed the summed population response over all recordings to all stimuli, and computed the Euclidian vector norm to each stimulus orientation. We defined a metric for symmetry as one minus the standard deviation of these vector norms divided by their average. A perfect symmetry of 100% would therefore mean that the vector norm of all orientation responses were identical. A symmetry of 0% would be the result if the standard deviation over orientation responses were as large as their mean. Using this metric, we found the orientation manifold to be 95.6% symmetric (fig. 5e), which – in our opinion – qualifies as exceptionally high for a noisy empirical data set. Even when computing this metric for individual recordings, we found very high symmetry values, ranging from 90-96% (fig. 5f). Finally, we decomposed all population responses into two parts: their gain-aligned component and the component in any other, orthogonal direction. This simple analysis also showed in a very direct way that neural noise is predominantly gain-aligned: on average across all n = 11 recordings, 69% of all neural variability lay along the 1-dimensional gain-axis (t-test of gain-aligned versus other noise, p = 2.4 x 10^−6^) (fig. 5g,h). This means that the remaining 23 – 159 dimensional subspace (depending on the size of the recording’s neuronal population) only contained a third of the neural variability, despite being many times larger, hence confirming the plausibility of manifolds as gain-scalable entities (fig. 5i).

Thus far our results provide a novel perspective on neural noise at multiple timescales, and we have shown that the bulk of this noise is restricted to a population gain axis that does not impair neural coding efficiency. While we recorded our stimuli without a task component, we did record the pupil size of our mice. Since pupil size is strongly correlated with arousal, we therefore wondered if population gain might be modulated by the mouse’s level of arousal. We calculated the population gain and pupil size on a per-trial basis and indeed found a positive correlation (fig. 5j, Pearson r = 0.15, p = 2.9 x 10^−2^), confirming our hypothesis that the population gain may be related to behavioural arousal.

## Discussion

We found that neuronal population responses are excessively variable, both at short (1 – 100 ms) and long timescales (0.1 – 10 s). At a subsecond timescale, neural dynamics are dominated by intermittent flares of activity on an otherwise relatively quiescent background (fig. 1). These dynamics are reminiscent of up/down states that have previously been reported in the membrane voltage of single cells^39–43^. We studied these flares in more detail and found that they were layer-specific: dense activation of supragranular cells appeared to drive the formation of these flares, while cells in deeper cortical layers showed less variable spiking rates (fig. 2). Interestingly, these flares also appeared in a large-scale computational model of the early visual system, where they arise from local network interactions (fig. 3). Model simulations suggest that spatiotemporally localized V1 flares may differentially affect subcircuits in V2, by driving strongly connected cells and (indirectly) inhibiting weakly connected cells in downstream cortical areas (fig. 3g-i). At longer timescales, on the order of second-long trials, the temporal coordination of spiking activity between cells gives rise to noise correlations. We found that population activity at these timescales obey a linear neural scaling law: an increase in average activity is accompanied by an equally strong increase in the standard deviation of neural activity (fig. 4). Moreover, we found that the bulk of all neural variability is aligned to a single gain axis (fig. 5). This model for neural noise describes the distribution of spiking rates across trials with stunning accuracy (R^2^ = 0.994), whereas a multivariate Gaussian model performs mediocrely (R^2^ = 0.517). These results led us to formulate a modified model for how neuronal populations encode information. This model incorporates the main gain-aligned noise mode as an invariant subspace (fig. 5i), effectively making the feature manifold a gain-scalable entity. Finally, we have shown that neural gain, like average spiking rate, may be a behaviourally-relevant variable that correlates with an animal’s pupil size; a common proxy for arousal^50–52^ (fig. 5j). While populations of neurons might appear excessively noisy, we found that population noise obeys strict mathematical structures that could make neural variability functional and prevents it from hampering feature coding. Mechanistically, population variability may serve as an inter-areal gating mechanism; computationally, it allows invariance to the dominant mode of noise.

One limitation of this study is that it is currently unclear whether the results we found in primary visual cortex of mice generalize to other brain areas and species. Flares and single-cell up/down states share many characteristics, including temporal dynamics^53^. We therefore believe it is likely that the flares we report here are the population-level result of the synchronization of up/down states between cells in a local circuit. If this is true, it would suggest that gain-dominant noise and flare dynamics are ubiquitous across cortex, as up/down states have been reported in different brain areas and animal species^39,41,42^. Flare-dynamics are also similar in many ways to the packets of activity previously found in auditory cortex^54,55^ – perhaps these phenomena are even different descriptions of the same underlying process. These auditory packets occur on a similar timescale of 10 – 100 ms, are hypothesized to be involved in gating signals, and are also characterized by transient periods of higher activity on a low-activity baseline^54,55^. Whereas we expect flare dynamics to generalize across cortex, different processes may dominate in subcortical structures, such as the superior colliculus and the lateral geniculate nucleus, where neural responses are more event-driven than in visual cortex^56–58^. Future studies will have to be performed to test these predictions.

Another open question is the relationship between neural oscillations and flares. We investigated whether we could detect a rhythmic signature of the occurrence of population flares, but they appeared to be occurring randomly – an autocorrelogram of flare occurrence showed only a long multi-second increase in flare probability with no evidence of a subsecond frequency at which their recurrence would be increased (data not shown). Linking the occurrence of flares to neural oscillations was not our main aim, however, so it is possible that more refined analyses of larger data sets could still reveal a connection. The relatively long 10-100 ms timescale of flares themselves indicate that if any connection between neural oscillations and flares exists, it is more likely to be in the alpha-oscillation range (∼10 Hz) than in the gamma-oscillation range (∼50 Hz).

A link between flares and divisive normalization seems an interesting possibility, however. Divisive normalization is a neural computation that rescales the overall level of population activity to a homeostatic baseline^59^. In doing so, it generates a winner-takes-all effect that has been proposed to underlie attentional selection^49,60–62^. The involvement of interneurons in the flare/quiescent dynamic we found in real data as well as in our model, is reminiscent of pyramidal/interneuron interactions hypothesized to underlie divisive normalization, and it is possible that flares are a side-effect of this neural computation. A slight offset in the peak latency of excitatory and inhibitory activity would be sufficient to generate the population flares we have observed. We therefore hypothesize that the implementation of divisive normalization in mouse V1 is such that this process is driven primarily by cortical feedforward output layers 2/3, and its function is indeed not only to normalize the neural response, but also to allow selective gating of neural signals at 10 – 100 ms timescales. Moreover, since our shuffling procedure showed that excess variability was undetectable at timescales under 1 ms (fig. 4c), this also puts a lower bound at the duration of this computation.

This study also connects short timescale operations, such as divisive normalization, to longer timescale phenomena, such as trial-level noise correlations. Both are the result of the temporal coordination of spiking activity between cells, but simply at different timescales. Population activity at the level of one-second trials revealed a linear neural scaling law, where an increase in average activity is accompanied by an equally strong increase in standard deviation. Note that this is not what is expected from cells that spike like Poisson process neurons – Poisson processes are linear in variance over the mean rather than standard deviation over the mean. Spiking correlations are therefore structured in very particular way that gives rise to an information constancy, invariant to the gain of the population activity (eq. 12-16). This relationship is so specific and unexpected (fig. 4e; data linearity = 0.92 versus Poisson linearity = 0.50), and it explained our data so well (fig. 4j, r = 0.89, p = 4.7 x 10^−18^) that we call it a “neural scaling law”. This does not mean we believe it governs neural activity as strictly as a true mathematical law would; it simply indicates a strong and non-trivial relationship.

The idea that the population gain is a fundamental mode in neural circuits is also supported by the accuracy of our gain-model for neural noise (fig. 5a-d). Despite requiring fewer parameters than the Gaussian model for neural noise (*n + 2* versus *2n*), our gain-model showed an almost un-neuroscientific level of predictability (R^2^ = 0.994) of the distribution of neuronal population noise. In contrast, the very common Gaussian model for neural noise performed rather poorly (R^2^ = 0.517). An obvious counterargument would be that we enforced independence between neurons, an assumption we know is violated in the brain. However, fitting a full multivariate Gaussian to the data would require *O(n*^2^) parameters, as the covariance matrix contains all pairwise interaction terms between neurons. Moreover, a further improvement in predictability is unlikely, since the gain model already achieved almost perfect prediction (R^2^ = 0.994). Therefore, even if a full a log-mvn model would predict the neural data perfectly, the gain-noise model would still be the more parsimonious description.

The combination of a linear neural scaling law with a gain-dominant mode of noise, and the observation that the orientation tuning manifold in V1 is highly symmetric (95.6% symmetry), inspired us to formulate a modified model for how neuronal populations encode information. In this model, the encoding subspace consists of an orientation manifold that multiplicatively scales with the population gain. This model stands in contrast to the classical model, where the population gain is simply one of many possible dimensions for neural noise. The main difference is therefore that we propose that the population gain is distinct from other (noise) dimensions in neural space. It is not only the dominant noise mode, making up 99.4% of all inter-trial variability in population rate (fig. 5c), but also lies in the nullspace of the encoding manifold. This conclusion can be drawn from the fact that the orientation manifold is 95.6% symmetric in terms of vector norms (fig. 5e,f). This means that only a maximum of 4.4% of the manifold is aligned to the population gain axis, and it is therefore effectively orthogonal to the population gain axis. The advantage of our gain-invariant neural coding model is therefore three-fold: 1) it needs fewer free parameters than the alternative model of a multivariate-gaussian noise plus ring manifold, 2) our model describes the data exceptionally well, and 3) it provides an explanation as to why neural systems can function despite showing an excessive amount of variability: the structure of those neural fluctuations generates invariance to the dominant mode of noise, which is aligned to the population gain axis.

## Methods

### Neuropixels data collection, preprocessing and selection

The electrophysiological data used here are previously described elsewhere^63^. In short, we performed 21 repeated-insertion recordings with Neuropixels across various brain regions in seven C57BL/6J (Janvier) mice. All mice were housed in a 12 h/12 h dark/light cycle with ad libitum access to food and water and were awake during recording. Spikes were sorted post-hoc using Kilosort^64^, and electrode location was determined by aligning histological slices and neurophysiological landmarks to the AllenCCF mouse brain atlas (https://github.com/cortex-lab/allenCCF) using the UniversalProbeFinder^65^. Of these 21 recordings, 16 penetrated primary visual cortex (V1) and were included in the initial data selection. The recording setup presented 40 repetitions of drifting gratings of 24 directions (spaced in 15-degree steps) and was controlled using Acquipix^66^. Since the detection of population-level phenomena requires a sufficient number of neurons and spikes, we discarded all recordings where the average number of spikes per drifting grating trial summed over all neurons was < 90. The remaining 11 data sets contained on average 64.1 V1 cells per recording, and 5.2 spikes per second per neuron during visual stimulation. We did not select for stimulus-responsiveness and/or tuning, and included all cells with an average firing rate of at least 0.1 Hz across the entire recording. All experiments were approved by the animal ethics committee of the Royal Netherlands Academy of Arts and Sciences, in compliance with all relevant ethical regulations.

### Decoding analyses

All decoding analyses used a multiclass logistic regression decoder with mild regularization (*λ=1*) and leave-one-repetition-out cross-validation^67^. The function *doCrossValidatedDecodingLR* that implements this algorithm can be downloaded here: https://github.com/JorritMontijn/DecodingAndInformation. We defined the “decoder confidence” we report throughout the manuscript as the posterior probability the decoder assigns to the correct stimulus. Using confidence instead of accuracy has the benefit that single-trial (or single spike-block) estimates are scalars rather than binary decisions. Note that accuracy is generally higher than confidence, as any confidence above 1/*n*, where *n* is the number of stimulus classes, may be sufficient to reach a “correct” decision.

### Spike-block analyses

Most analyses in neurophysiology are performed on neural activity averaged over some time window; for example 25 ms bins in a peri-stimulus time histogram (PSTH) analysis. This fixes the time window, and allows for a variable number of spikes per bin. This approach is versatile, but a big drawback is that many analyses are biased by the number of spikes, and this approach is therefore unsuited to comparing high-activity epochs with low-activity epochs. We therefore designed an alternative spike-block approach. In this analysis, we fixed the number of spikes per bin and instead let the time window size vary. For each recording, we chose the spike-block size to be a fraction of the average number of spikes during a single trial summed over all neurons in the population. Let ***A*** be a *[p* by *q]* matrix of firing rates for all *p* trials and *q* neurons, and ***a*** the *[p* by *1]* vector of population firing summed over all neurons. Then the spike-block size *l* is

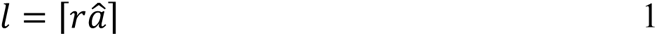

where *â* is the mean of ***a***, and ⌈ ⌉ is the ceiling function. We chose the value of *r* to obtain a good balance between the number of spikes per block and the number of spike blocks. All analyses presented throughout the paper use *r* = 1/30, which equates to an average block duration of 33 ms. We repeated our analyses for *r* = 1/20 and found that our results are not dependent on the exact choice of *r*.

For each trial, we split the spikes of all neurons during that trial into equally sized sets *b*, starting with the first spike after stimulus onset. For example, if ***v***^1^ is a sorted vector of spike times for trial 1, and the spike block size *l* is 15, then spike block 2 (*b_2_*) is the set of spikes ***v****^1^_16…30_*. More generally, for trial *t*, block number *j*, block size *l*, and spike number *i*:

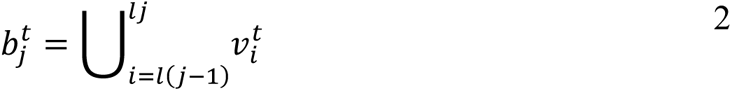

Note that here we use superscript to indicate trial number and not exponentiation. If fewer than *l* spikes remain at the end of a trial, so the last block cannot be completely filled, we discard the last incomplete block. We treated each spike block as an independent “trial” for all further analyses; for example, using spike blocks as trials allowed us to decode the stimulus information over time (Fig. 2c). Note that we show the instantaneous population firing rate for illustrative purposes (e.g., Fig. 2b), but the spike block’s firing rate is defined independently of this instantaneous rate. We instead define the firing rate of a spike block as the inverse of the block’s duration, as each block has the same number of spikes: i.e., 15 spikes in 15 ms equals 1000 Hz, whereas 15 spikes in 30 ms equals 500 Hz.

### Shuffling and other data randomization procedures

We randomized the real neural data in multiple ways to investigate the effect of removing specific properties (e.g., fig. 4c). For example, to test the effect of flares of neuronal population activity, we compared real data to those where we permuted the spike times across trials (“Shuffled”, see fig. 1j). Descriptions of each procedure will follow hereafter.

#### Shuffled

*Shuffled* data was created independently for each neuron and stimulus class. Let ***x*** and ***t*** be n-element vectors, where *n* is the number of spikes fired by neuron *k* over all *j=1…p* repetitions of stimulus class *k*. *x_i_* is then the stimulus repetition number during which spike *i* was fired, while *t_i_* is the latency of spike *i* relative of the onset of the stimulus. Finally, ***τ*** is the p-element vector of onset times corresponding to the repetition numbers in ***x***; so max(***x***) = *p*. The overall spike time *v_i_* is therefore given by:

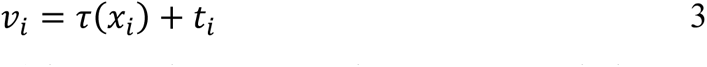

Now we permute ***x*** (without replacement) by a random permutation vector ***π*** such that

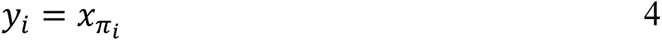

This means that each spike gets assigned to a different trial, but in such a way that the number of spikes per trial remains unchanged, since the number of entries per trial in ***x*** and ***y*** remain the same:

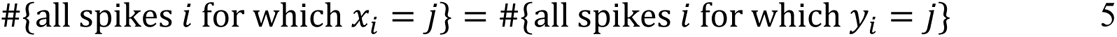

Where # is the set cardinality. Now we create a shuffled spike time vector ***w*** using these permuted trials:

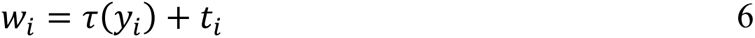

Note that this procedure preserves not only the trial-level spike counts, but also the trial-averaged dynamics per stimulus class. ***t*** is unaltered, so shuffled data ***w*** still shows an onset peak, for example. This procedure only removes short-term temporal coordination across neurons, allowing us to investigate specifically the presence of population flares.

#### Poisson

*Poisson* data was created as a null-hypothesis baseline, where we generated spike times for each neuron *k* by sampling inter-spike intervals from an exponential distribution P associated with the stationary Poisson point process that we matched individually to each neuron’s overall spiking rate *λ_k_* during stimulus presentation over all trials. For neuron *k*, the *Poisson* spike time data *w_i_* for spike number *i* is therefore

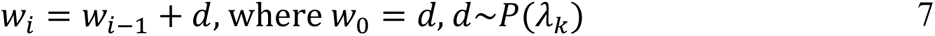

This procedure destroys all flares, trial structure, noise correlations, and stimulus information.

#### Poisson x Gain

An intermediate step between the aforementioned *Poisson* data and real data is a gain-modulated Poisson process. As above, we generate random spike times for each neuron, but we also modulate the rate parameter *λ* with a trial-based population-gain *γ* we calculated from the real neural data. As above, let ***A*** be the matrix of firing rates for all trials and neurons, ***a*** the vector of population firing summed over all neurons, and *â* the average population rate over trials. We then define the population gain *γ* for trial *j* as

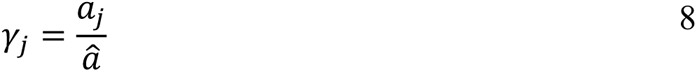

The gain-modulated Poisson-process spike time *w_i_* for spike number *i* during trial *j* is then

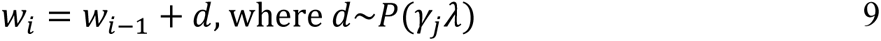

While similar to the unmodulated *Poisson* data, the comparison of the *Poisson x Gain* data to the aforementioned Shuffled data allowed us to investigate to what extent any difference between them was explainable by trial-level gain fluctuations on the one hand (present in *Poisson x Gain*), and stereotyped within-trial dynamics and stimulus tuning differences on the other (absent from *Poisson x Gain*).

#### Uniform

We investigated the effect of trial structure (e.g., the presence of an onset peak) by comparing *Shuffled* data to data obtained from a procedure where we enforced a uniform spiking probability over each trial, for each neuron independently. Using the same convention as above, uniform spike times were generated as follows:

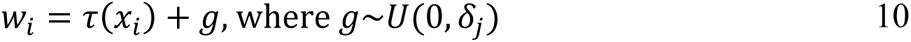

Here, *δ_j_* is the duration of trial *j*, and *U*(0*, δ_j_*) is the uniform probability density over the interval (0, *δ_j_*). Note that this procedures assigns random spike latencies, but – like *Shuffled* data – keeps the number of spikes per trial unchanged.

#### Rnd SpT

The random spike time (*Rnd SpT*) data is similar to *Shuffled* data, except here we also randomized the number of spikes per trial by randomly permuting ***τ*** rather than ***x***. Since we performed this procedure for each stimulus class independently, however, the average tuning to different stimulus classes remains intact. This procedure allowed us to investigate the effect of trial-level noise correlations (when comparing with *Shuffled*), and to investigate the effect of stimulus tuning (when compared with *Rnd xStim*; see below). that since this procedure also relies on permuting rather than random sampling, the trial structure remains intact. Using the same conventions as equation 7, spike times for *Rnd SpT* are defined as:

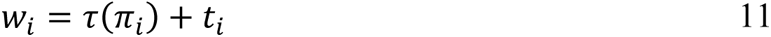

Note that ***π*** is now an *n*-element vector (*n* being the number of spikes of one neuron for one stimulus orientation), as each spike is independently assigned a trial at random.

#### Rnd xStim

Finally, we created data where we only kept the trial structure and overall firing rate per neuron, but destroyed all other properties. In this random spike time across stimuli (*Rnd xStim*) data, we altered spike times like in equation 11, except we defined a single vector ***τ*** and a single vector ***t*** for all trials of all stimulus classes combined. Compared with *Rnd SpT*, this therefore also discards stimulus tuning.

### Description of computational model for multi-area simulations

Our large-scale computational model has been described in a pre-print, but we will include its description here as well for completeness^47^. The model consists of a retina, LGN, V1 and V2, based on descriptions of the early visual system in the macaque. The units in the retina and LGN are uncoupled point processes: the retina is a filter-based 2D activity map with *iid* Gaussian noise on top of the filter response, and the LGN turns this activity map into spikes using a stochastic firing probability. The LGN stage feeds into the first fully simulated cortical stage (V1), using a connectivity pattern that gives rise to Gabor-like receptive fields in V1, similar to^16^ and ^68^. Excitatory V1 cells then project to V2 to create angle and texture-like receptive fields in V2 cells. All cortical cells are simulated using a conductance-based leaky-integrate-and-fire model. Cells in each cortical area (V1 and V2) are sparsely recurrently connected. A full description of the model can be found in the supplemental methods.

### Analysis of sd/mean linearity at long timescales

We analyzed the linearity of standard deviation over the mean over spike counts per bin (fig. 4d). We chose timescales from 100 ms up to 10 s – the range where population flares had negligible effect on the population variability (fig. 4c) – and for each timescale, we calculated the spike count per bin. Therefore each timescale yielded a standard deviation and mean, and to these points we fitted a line. We obtained an explained variance of the linear fit (R^2^) for each recording, and plotted the mean ± standard error over recordings (n=11) in fig. 4e. We repeated the analysis for different data-randomizations (see above).

### Proof of gain invariance with constant coefficient of variation

The coefficient of variation is defined as the standard deviation divided by the mean. A simple derivation can show that under the assumption of Gaussian distributions with equal variance, the neural information, which we define here as the inverse of the misclassification error, is constant regardless of scaling of the average activity. With a constant coefficient of variation *c*, the neural gain *γ* will apply equally to the mean *μ* and standard deviation *σ* of the neural response:

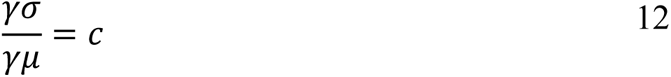

In this scenario, two stimulus orientations *θ1* and *θ2* are encoded by a neuronal population, where the distribution of firing rates across the decision axis follows Gaussian distributions *g1* and *g2* for the two stimulus orientations. With means *γμ_g1_* and *γμ_g2_* and equal variance *γσ*^2^, the optimal decision boundary *d* is therefore

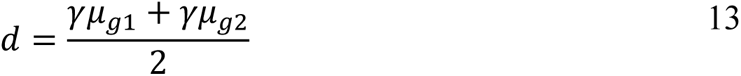

The misclassification error *ε* for *g1*, with *μ_g1_* < *μ_g2_*, is

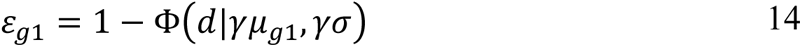

Where Φ is the cumulative normal distribution. With two distributions of equal variance, we can freely choose our point of origin without loss of generality. Choosing our origin such that *γμ_g1_* = –*γμ_g2_*, we obtain a symmetry where *d = 0*. We can then rewrite the above equation as

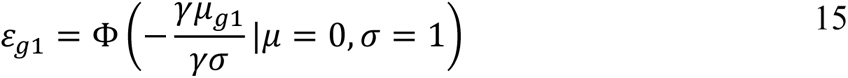

Since *g1* and *g2* are symmetric, *ε_g1_* = *ε_g2_*. With a constant coefficient of variation *c*, where *c = γσ/γμ*, the total misclassification error is:

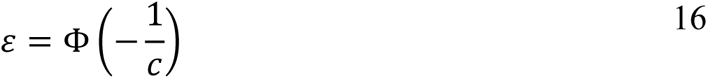

With a constant coefficient of variation *c*, the neural information 1/ε is therefore gain-invariant.

### Procedure to enforce sd/mean scaling data

As before, we define ***A*** to be a *[p* by *q]* matrix of trial-average firing rates for all *p* trials and *q* neurons. Averaging over neurons yields a *[p* by *1]* vector of average population firing rates ***m***, and taking the standard deviation (sd) over neurons yields a *[p* by *1]* vector ***s*** of the sd of firing rates within the population for all *j = 1*…*p* trials. For each trial *j*, we subtract the population average from matrix ***A***:

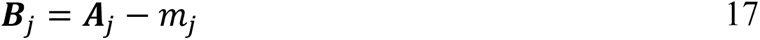

The population mean for each trial in ***B*** is therefore 0, as all trials now have a population rate of 0. However, note that the sd over neurons for each trial is unchanged. Now we enforce a uniform sd for each trial:

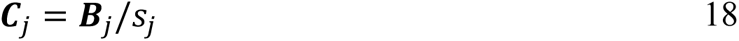

Matrix ***C*** is therefore z-scored per trial, with each trial having a population mean rate of 0 and a population sd of 1. Now we will enforce a linear sd/mean relationship by multiplying each trial in matrix ***C*** by its original population rate and adding back the original mean rate:

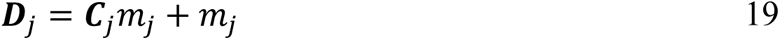

This procedure enforces that the coefficient of variation is exactly 1 for the sd/mean computed over all neurons for each trial.

### Gaussian noise and gain noise models

We investigated two models for neural noise: (log-)normal (aka, Gaussian) and gain-based. Gaussian noise models are commonly used in neurophysiology, and come in two flavours: linear and logarithmic. Linear Gaussian models are easier to work with and are more common, but since neuronal firing rates cannot be negative, logarithmic Gaussian models may be more accurate. In this analysis, we investigated how well the distribution of population firing rates, as well as of single neuron firing rates, is described by the model. First, we create matrix ***A***, a *[p* by *q]* matrix of firing rates for all *p* bins and *q* neurons, and ***a*** the *[p* by *1]* vector of total population rate per bin, summed over all neurons. We repeat this procedure for five timescales, where we set the bin width to be 10, 50, 100, 500, or 1000 ms. For each timescale, we perform the following procedure. First we calculate the mean ***μ*** and standard deviation ***s*** over bins for each neuron, and use the standard deviations to construct a diagonal covariance matrix, with *j* and *k* indexing over neurons:

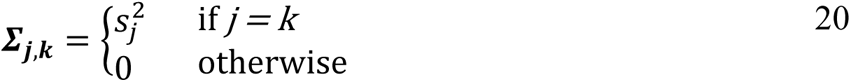

Then we create predicted matrix ***B*** by taking *p* random samples from this multivariate Gaussian:

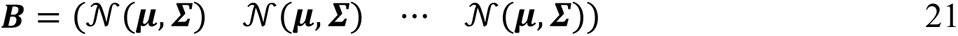

Now we can quantify how well this model predicts the distribution of population firing rates by comparing vectors ***a*** and ***b***, the population rate per bin, summed over all neurons for matrices ***A*** and ***B*** respectively. First we sort the values in ***a*** and in ***b***, and then compute the explained variance between them:

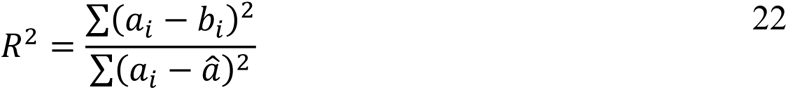

As before, *â* is the mean over ***a***. We calculate the accuracy of single-neuron predictions by constructing sorted vectors of all elements of ***A*** and ***B***, and then computing R^2^ as described above. For the log-normal noise model, we substitute the multivariate normal distribution in eq. 21 by a multivariate log-normal distribution.

Next, we constructed a gain-based neural noise model. While the two Gaussian models described above use 2 *q* parameters (mean and sd per neuron), the gain model uses only *q* + 2 parameters: a gain-axis ***v***, defined by the mean response of each neuron (so identical to ***μ***), and a mean *m* and sd *s* of the log-normal distribution of the gain distribution ***γ***. Like above, we then take *p* random samples from this model, with each sample *i* defined as:

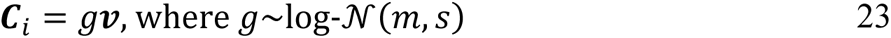

Finally, we also used a model where we defined a separate gain axis for each stimulus class. This procedure works as above, except we use a distinct gain vector ***v*** for each stimulus class.

## Supporting information

Supplementary Model Description

## Acknowledgements

We thank Valentina Riguccini for contributing to previously published data sets that are used in this study. This work was funded by a Royal Netherlands Academy of Arts and Sciences (KNAW) Fonds KNAW-Instituten grant. The authors declare no competing financial interests.

