## Supplementary Model Description for "Structured neural fluctuations can generate noise invariance and inter-areal gating at distinct timescales"

**Description for a large-scale computational model of the early visual system**

The following model has been previously described in a bioRxiv pre-print[1]. Still, for methodological completeness we will describe the model in full in the following paragraphs. Code to run this model is available online at https://github.com/JorritMontijn/SimulationsEVS.

**Filter-based input layers: Retina, LGN**

The retina layer in our model consists of two separate streams (ON-centre and OFF- centre) that are arranged in arrays of 32 x 32, with a spacing of 0.2 degrees visual angle between cells. For a patch of 6.4 x 6.4 degrees we simulated a visual input twice the size (12.8 x 12.8) to avoid edge effects. The rate-based activation in the retina for a cell at location (*x,y*) for ON-centre and OFF- centre streams were

$$R_{ON}(x,y,t) = [r_{baseline} + \boldsymbol{I} * \kappa(x,y,t|K_c,\sigma_c) - \boldsymbol{I} * \kappa(x,y,t-\delta|K_s,\sigma_s)]^+, \qquad 1$$

$$R_{OFF}(x,y,t) = [r_{baseline} - \boldsymbol{I} * \kappa(x,y,t|K_c,\sigma_c) + \boldsymbol{I} * \kappa(x,y,t-\delta|K_s,\sigma_s)]^+. \qquad 2$$

Here, $[\cdot]^+=\max(\cdot,0)$ denotes rectification, $\boldsymbol{I}$ is the input image, which depends on space and time, * denotes convolution, *c* indicates centre response, *s* indicates surround response and δ is a 3 ms delay between centre and surround responses. The spatiotemporal kernel $\kappa$ consists of a circularly symmetric Gaussian spatial profile and an exponential temporal impulse function,

$$\kappa(x,y,t|K,\sigma) = \frac{1}{\tau}e^{-\frac{t}{\tau}}\frac{K}{2\pi\sigma^2}e^{-\frac{x^2+y^2}{2\sigma^2}} \qquad 3$$

We truncated the filter for locations more than four times the standard deviation of the Gaussian profile away from its mean for computational efficiency. We used the parameters taken from [2] (see Table 1). The LGN layer simply transforms the $R_{ON}$ and $R_{OFF}$ analogue activity levels into stochastic spiking events, with an average rate of 40 Hz at full stimulus contrast.

**LGN-V1 connectivity structure: Gabor fields**

Each V1 cell has a Gabor-like receptive field, with several parameters that varied across cells (see Table 1): phase ($\psi$), orientation ($\theta$), spatial frequency ($1/\lambda$), spatial length ($\sigma_y$), spatial width ($\sigma_x$), horizontal centre ($\mu_x$), and vertical centre ($\mu_y$). Given these parameters, we created a Gabor patch *G*,

$$G(x, y, \mu_x, \mu_y, \sigma, \lambda, \theta, \psi) = exp\left(-\frac{(x-\mu_x)^2 + \left(y-\mu_y\right)^2}{2\sigma^2}\right) cos\left(\left(\frac{2\pi}{\lambda}\right)\left((x-\mu_x)\, cos\,\theta + \left(y-\mu_y\right) sin\,\theta\right) + \psi\right). \quad 4$$

With each V1 cell assigned a Gabor field, we next calculated the connection probability weight for ON-centre cells at retinoptic locations (*x, y*) from LGN to a V1 cell *i* as follows,

$$r(x, y)_{V1,LGN} = \frac{[G_i(x, y)]^+}{\sum_{x,y}[G_i(x, y)]^+}. \quad 5$$

For OFF-centre cells, we used the same formula, but inverted the sign of *G*. We then connected the required number of LGN cells from each stream to a V1 cell by choosing an LGN cell to project with a probability proportional to its weight described above, relative to that of other cells. If a connection was made, we assigned to each a weight equal to its corresponding probability, *r(x,y)*, a conductance  and a Gaussian delay (see Table 1 for parameter values).

**Leaky-integrate-and-fire (LIF) layers V1 and V2**

For the fully simulated cortical layers, we used a conductance-based leaky-integrate-and-fire model of pyramidal cells and interneurons, as described in [3] and originally in [2]. We simulated 1200 V1 cells and 1200 V2 cells, of which 80% were excitatory and 20% were inhibitory. For post-synaptic cell *i*, the membrane voltage, $V_i$, evolves according to

$$C_i \frac{dV_i(t)}{dt} = -g_i^{Exc}(V_i(t) - V_i^{Exc}) - g_i^{Inh}\big(V_i(t) - V_i^{Inh}\big) - g_i^{Leak}\big(V_i(t) - V_i^{Leak}\big) - g_i^{AHP}\big(V_i(t) - V_i^{AHP}\big). \quad 6$$

Here, $g^{Exc}$ denotes the total excitatory post-synaptic conductance due to the feedforward and recurrent input, $g^{Inh}$ similarly denotes the total inhibitory conductance, $g^{Leak}$ denotes the passive leak conductance, and $g^{AHP}$ the after-hyperpolarization conductance due to prior spiking. $V^{Exc}$, $V^{Inh}$, $V^{Leak}$ and $V^{AHP}$ denote the reversal potentials for excitatory, inhibitory, leak and AHP inputs respectively. $C_i$ the membrane capacitance for neuron $i$. Whenever a cell's membrane voltage exceeded -55 mV, we used a delta function to simulate spiking behaviour, and reset the cell's membrane voltage to its resting potential ($V^{Leak}$) on the next time step. As the simulation time step was 0.5 ms, our cells had an absolute refractory period of 1 ms. We used the following (approximately) biologically plausible parameters for all cells; see Table 1, which summarises the parameters.

**Cortical recurrent connectivity: similarity- & locality-based**

Cortical connectivity was determined on the basis of receptive-field similarity, where the weighting of the similarity is controlled by a locality hyperparameter $\beta$. $\beta$ = -1 corresponds to a uniform connectivity (i.e., ignoring similarity), and $\beta$ = 0 corresponds to a proportional weighting, as described above for LGN-V1 connectivity. We set $\beta$ = 0 for excitatory neurons, and $\beta$ = -0.5 for inhibitory neurons. This means that inhibitory neurons project more uniformly to less similarly tuned cells than excitatory neurons, as has been shown in mouse visual cortex [4]. We calculated an element-wise similarity $\rho$ in receptive fields of cells $i,j$ as follows. First, we normalised each receptive field **G** to be mean-zero:

$$\boldsymbol{G}' = \boldsymbol{G} - \iint \boldsymbol{G}\,. \quad 7$$

The similarity metric $\rho$ (in the range [-1 1]) is given by

$$\rho_{i,j} = \frac{\boldsymbol{G}'^{T}_{i}\boldsymbol{G}'_{j}}{\sqrt{\boldsymbol{G}'^{T}_{i}\boldsymbol{G}'_{i}\boldsymbol{G}'^{T}_{j}\boldsymbol{G}'_{j}}}. \quad 8$$

Here, $^T$ indicates transpose. We rectified all negative $\rho$ values to 0, and created a cumulative probability vector **r** based on $[\rho]^+$, as described above for LGN-V1 connectivity. For a number of connections $n$ originating from a source neuron, we randomly chose $n_{proportional} = n*(1-|\beta|)$, rounded to the nearest integer, connections based on **r**. The remaining $n_{non\text{-}proportional} = n - n_{proportional}$ connections were chosen either according to a uniform random distribution across neurons when $\beta < 0$. This way, connectivity is a mixture of uniform and $[\rho]^+$-proportional connections for $-1 < \beta < 0$.

**Visual stimulus parameters**

Visual stimuli were always centred in the middle of the simulated patch of visual space and consisted of drifting sinusoidal gratings within a circular aperture 5° in diameter, with a cosine-ramped edge ($\cos^2(r/\lambda_0)$ where $\lambda_0 = 2°$ and $r$ is distance from the edge of the 5° diameter patch, measured in degrees and ranging from 0 to π degrees) leading to a neutral-grey background. Unless stated otherwise, all stimuli had a pre-stimulus uniform neutral-grey blank period of 100 ms, after which the stimulus was presented for 500 ms, and ended with another 100 ms blank period. We used a spatial frequency of 0.25 cycles per degree, a temporal frequency of 2 Hz, full contrast and luminance, and a random starting phase.

**Programmatic implementation**

We increased computational efficiency of our model by vectorising the calculation of spike-induced post-synaptic potentials across spike history and synaptic connections, thereby requiring at each time step only a for-loop across cells, rather than across synaptic connections. This is difficult to achieve in a setup such as ours, where heterogeneous synaptic delays and exponentially decaying synaptic conductances act on a single target neuron, because calculating the resulting action on the membrane voltage requires incorporating all heterogeneous pre-synaptic spike times from various time steps in the past. We achieved this vectorization by using a sparse dynamic cylindrical array to store recent spiking activity for all neurons in a reusable buffer variable, as described originally by [5]. The algorithm (see below for a pseudo-code description) uses a vector $\boldsymbol{\nu}$ that stores the number of recently emitted spikes per cell, a vector $\boldsymbol{\pi}$ that keeps track of the current position on the cylindrical array for each cell, and a cylindrical array **T** of size [cells x spikes] that stores the time stamps of recent spikes. Looping across cells, single-synapse inputs are stored in a vector, where each element $q$ corresponds to a single synaptic connection from cell $j$ to cell $i$. For each source cell $j$, the synaptic drive in all its output synapses can be calculated using vector operations. We loop through all neurons in $\boldsymbol{v}$, ignoring all cells that have zero recent spikes, yielding another performance boost. Let $n$ be a non-zero element for cell $j$ in $\mathbf{v}$, and $\mathbf{p}$ be an $n$-dimensional vector of indices computed from $\boldsymbol{\pi}$ that correspond to the locations in $\mathbf{T}$ where the most recent spikes are stored. Now the recent spike times of cell $j$ can be accessed simply through $\mathbf{T}(j,\mathbf{p})$, providing a vector $\boldsymbol{\tau}$ of time stamps. We next created a matrix $\boldsymbol{\Delta}$ of size [spikes x synapses] that contains the spiking time relative to the current simulation time $t$, normalised for the synaptic delays $\boldsymbol{\delta}$. For spike $k$ and synapse $l$, this relative spiking time can thus be written as

$$\Delta_{k,l} = t - \delta_k - \tau_l. \quad 9$$

Note that this operation is easily written as a vectorised calculation in any Matlab implementation. The sum total of excitatory conductances $\boldsymbol{s}$ for all target synapses of $j$ are now given by

$$\boldsymbol{s} = \boldsymbol{w} \cdot \boldsymbol{c} \cdot \boldsymbol{h}(\boldsymbol{\Delta}, \tau_j) \quad 10$$

Here, · indicates element-wise multiplication, $\boldsymbol{w}$ is a vector of synaptic weights, and $\boldsymbol{c}$ is a vector of synaptic conductances, $\tau_j$ denotes the synaptic time constant for cell $j$, which depends only on whether the cell is excitatory or inhibitory, and $\boldsymbol{h}$ is the exponential synapse function that outputs a sum total value for each synapse. As above, note that this function is completely vectorised over synapses, but for clarity we here write the function for a single synapse $l$,

$$h_l(\boldsymbol{\Delta}_l, \tau_j) = \sum_k \left( [\Delta_{k,l}]^+ \frac{1}{\tau_j} e^{1-\frac{\Delta_{k,l}}{\tau_j}} \right). \quad 11$$

Synaptic delay and conductance parameters are provided in their corresponding paragraphs detailing the connectivity structure. Finally, the resulting excitatory and inhibitory input per cell ($E^{Exc}$ and $E^{Inh}$) can be calculated by summing all synaptic inputs in **s** to their postsynaptic targets. This step is also easily vectorised, using the built-in Matlab function *accumarray()*. In Matlab pseudo-code, this algorithm can be written as follows:

```
%loop through all source neurons that have > 0 recent spiking events
for j=find(nu > 0)
    %get indexing vector of positions on cylindrical array
    p = getCycPosVec(pi(j), nu(j));

    %get vector of spike times from cylindrical array
    tau = T(j,p);

    %get indices of target synapses for this cell
    vecTargetSynapses = cellSynTargets{j};

    %get synaptic time constant for this cell
    tau_j = vecTauSyn(j);

    %get normalised spike time matrix
    Delta = bsxfun(@minus,t-delta(vecTargetSynapses),tau);

    %calculate PSPs
    s(vecTargetSynapses) = w(vecTargetSynapses) .* ...
        c(vecTargetSynapses) .* h(Delta, tau_j);
end
```

| **LGN network parameters** | | |
|---|---|---|
| **Symbol** | **Meaning** | **Value** |
| *N* | Number of LGN neurons | 2048 |
| *N (ON)* | Number of ON-centre cells (50%) | 1024 |
| *N (OFF)* | Number of OFF-centre cells (50%) | 1024 |
| *Nsyn (LGN-Exc)* | Number of LGN to E synaptic connections | 144 |
| *Nsyn (LGN-Inh)* | Number of LGN to I synaptic connections | 96 |
| *Gsyn (LGN-Exc)* | Conductance of LGN to E synapses | 0.264 |
| *Gsyn (LGN-Inh)* | Conductance of LGN to I synapses | 0.288 |
| *Dsyn (LGN-Exc)* | Synaptic delay between LGN and E cells (rectified to > 0) | i.i.d. random Gaussian |

| | | |
|---|---|---|
| | | (μ = 10 ms, σ = 7 ms) |
| *Dsyn (LGN-Inh)* | Synaptic delay between LGN and I cells (rectified to > 0) | i.i.d. random Gaussian<br>(μ = 5 ms, σ = 3 ms) |
| δ | Latency difference between centre and surround response | 3 ms |
| $\sigma_c$ | Spatial Gaussian width of centre | 0.176 |
| $\sigma_s$ | Spatial Gaussian width of surround | 0.53 |
| $K_c$ | Response magnitude constant of centre | 17 |
| $K_s$ | Response magnitude constant of surround | 16 |
| $\tau_c$ | Temporal exponential scale of centre | 10 |
| $\tau_s$ | Temporal exponential scale of surround | 20 |
| $r_{baseline}$ | Baseline firing rate | 15 |
| **Cortical network parameters** | | |
| **Symbol** | **Meaning** | **Value** |
| *N* | Number of LIF neurons per area (V1/V2) | 1200 |
| *N (Exc)* | Number of excitatory (E) V1 cells (80%) | 960 |
| *N (Inh)* | Number of inhibitory (I) V1 cells (20%) | 240 |
| *β (Exc)* | Locality hyperparameter for E cells | 0 |
| *β (Inh)* | Locality hyperparameter for I cells | -0.5 |
| *Nsyn (Exc-Exc)* | Number of E to E synaptic connections | 160 |
| *Nsyn (Exc-Inh)* | Number of E to I synaptic connections | 160 |
| *Nsyn (Inh-Exc)* | Number of I to E synaptic connections | 120 |
| *Nsyn (Inh-Inh)* | Number of I to I synaptic connections | 120 |

| | | |
|---|---|---|
| *Gsyn (Exc-Exc)* | Conductance of E to E synapses | 0.24 nS |
| *Gsyn (Exc-Inh)* | Conductance of E to I synapses | 0.32 nS |
| *Gsyn (Inh-Exc)* | Conductance of I to E synapses | 0.375 nS |
| *Gsyn (Inh-Inh)* | Conductance of I to I synapses | 0.25 nS |
| *Dsyn* | Synaptic delay between cortical cells (rectified to > 0) | i.i.d. random Gaussian ($\mu$ = 3 ms, $\sigma$ = 1 ms) |
| **Leaky integrate-and-fire neuron parameters** | | |
| **Symbol** | **Meaning** | **Value** |
| *dt* | Simulation time step | 0.5 ms |
| $V_0$ | Initial membrane voltage | i.i.d. random Gaussian ($\mu$ = -55.6 mV, $\sigma$ = 1 mV) |
| $V^{thresh}$ | Spiking threshold | -55 mV |
| $V^{Exc}$ | Reversal potential of excitatory channels | 0 mV |
| $V^{Inh}$ | Reversal potential of inhibitory channels | -70 mV |
| $V^{Leak}$ | Resting-state leak potential | -65 mV |
| $V^{AHP}$ | After-hyperpolarization (AHP) potential | -90 mV |
| $g^{Leak}$ *(Exc)* | Leak conductance excitatory cells | 25 nS |
| $g^{AHP}$ *(Exc)* | AHP conductance excitatory cells | 40 nS |
| *C (Exc)* | Membrane capacitance excitatory cells | 0.5 nF |
| $g^{Leak}$ *(Inh)* | Leak conductance inhibitory cells | 20 nS |
| $g^{AHP}$ *(Inh)* | AHP conductance inhibitory cells | 20 nS |

| | | |
|---|---|---|
| *C (Inh)* | Membrane capacitance inhibitory cells | 0.2 nF |
| **V1 tuning parameters** | | |
| **Symbol** | **Meaning** | **Value** |
| $\theta$ | Orientation | [0°, 3°, …, 177°] |
| $1/\lambda$ | Spatial frequency | [0.125°, 0.25°, 0.5°, 1°] |
| $\xi$ | Horizontal centre | i.i.d. uniform random in range [-3.2, 3.2] |
| $\upsilon$ | Vertical centre | i.i.d. uniform random in range [-3.2, 3.2] |
| $\sigma_y$ | Spatial length | 0.7° |
| $\sigma_x$ | Spatial width | 0.7° |
| $\psi$ | Phase | i.i.d. uniform random in range [0, 2π) |

*Table 1 Parameters used in the model.*
